# Pycea: a scverse toolkit for single-cell lineage tracing analysis

**DOI:** 10.64898/2026.10.30.763227

**Authors:** William N. Colgan, Eric Weine, Luke W. Koblan, Matthew G. Jones, Jonathan S. Weissman, Nir Yosef

## Abstract

Lineage tracing enables reconstruction of cellular histories in development, regeneration, and disease, but integrating lineage data into single-cell analysis workflows remains challenging. We introduce Pycea, a Python toolkit for lineage analysis and visualization, and TreeData, an extension of AnnData that stores lineage trees alongside cellular measurements. TreeData is compatible with Scanpy and other scverse tools, enabling integrated analysis without converting between formats. Pycea provides scalable methods for tree visualization, clade identification, distance comparisons, ancestral state inference, and lineage statistics. This software is open source and available at https://github.com/YosefLab/pycea.

## Introduction

Technological advances have dramatically increased the acquisition of single-cell lineage tracing data, with recently published studies including more than one million cells [1, 2]. These datasets combine measurements of individual cells with lineage trees describing their division histories, generated through live-cell tracking or molecular recording. In live-cell tracking, cell movements are directly observed using time-lapse microscopy, allowing lineage trees to be reconstructed from observed cell divisions [3–7]. In molecular recording, cells are engineered to accumulate heritable genomic marks that can subsequently be used to reconstruct their lineage relationships. These marks can be introduced using CRISPR–Cas9 [8–13], prime editing [14, 15], or other methods [16, 17]. Both approaches can pair lineage information with additional cellular measurements, including gene expression [2, 11, 18, 19], spatial coordinates [15], and cell morphology, enabling the study of cell states and behaviors across development, regeneration, and disease.

The scverse ecosystem has established a common framework for single-cell analysis in Python [20], centered on AnnData, a standardized data structure for storing single-cell measurements and associated metadata [21]. Extensions such as MuData and SpatialData add support for multimodal and spatial datasets [22, 23]. These interoperable data structures support a growing ecosystem of tools, including Scanpy [24] and Squidpy [25], allowing users to process, visualize, and analyze single-cell data without repeatedly converting between formats. However, the scverse ecosystem has so far lacked both a standardized data structure for storing lineage trees and a dedicated toolkit for lineage visualization and analysis. As a result, lineage information remains difficult to integrate into existing single-cell workflows.

Existing tools such as LinTIMaT, DCLEAR, Startle, TiDeTree, and LAML primarily focus on lineage tree reconstruction [26–31]. Our Cassiopeia package [32] additionally provides some plotting and analysis functionality, but integrating gene expression data is cumbersome because the `CassiopeiaTree` object does not natively store this modality or integrate with the scverse ecosystem. Similarly, users performing lineage analysis with phylogenetic tools such as ETE [33] need to manually transfer cell annotations, such as Leiden clusters, from AnnData into the ETE Tree object. Our goal is to eliminate this data juggling and enable seamless analysis of lineage trees alongside gene expression and other features.

Here, we introduce Pycea, a scverse-compatible toolkit for lineage analysis and visualization built around TreeData, an extension of AnnData that supports the storage of lineage trees along-side other single-cell measurements (Figure 1). TreeData serves as a drop-in replacement for Ann-Data, retaining compatibility with Scanpy and other scverse tools while enabling integrated lineage analysis. Pycea provides a suite of intuitive and scalable methods for visualizing lineage trees, identifying clades, comparing lineage distances, inferring ancestral states, and more. We demonstrate these capabilities using lineage tracing datasets from mouse development and a metastatic tumor model [2, 15]. Additional examples are available in the Pycea documentation.

**Figure 1:**
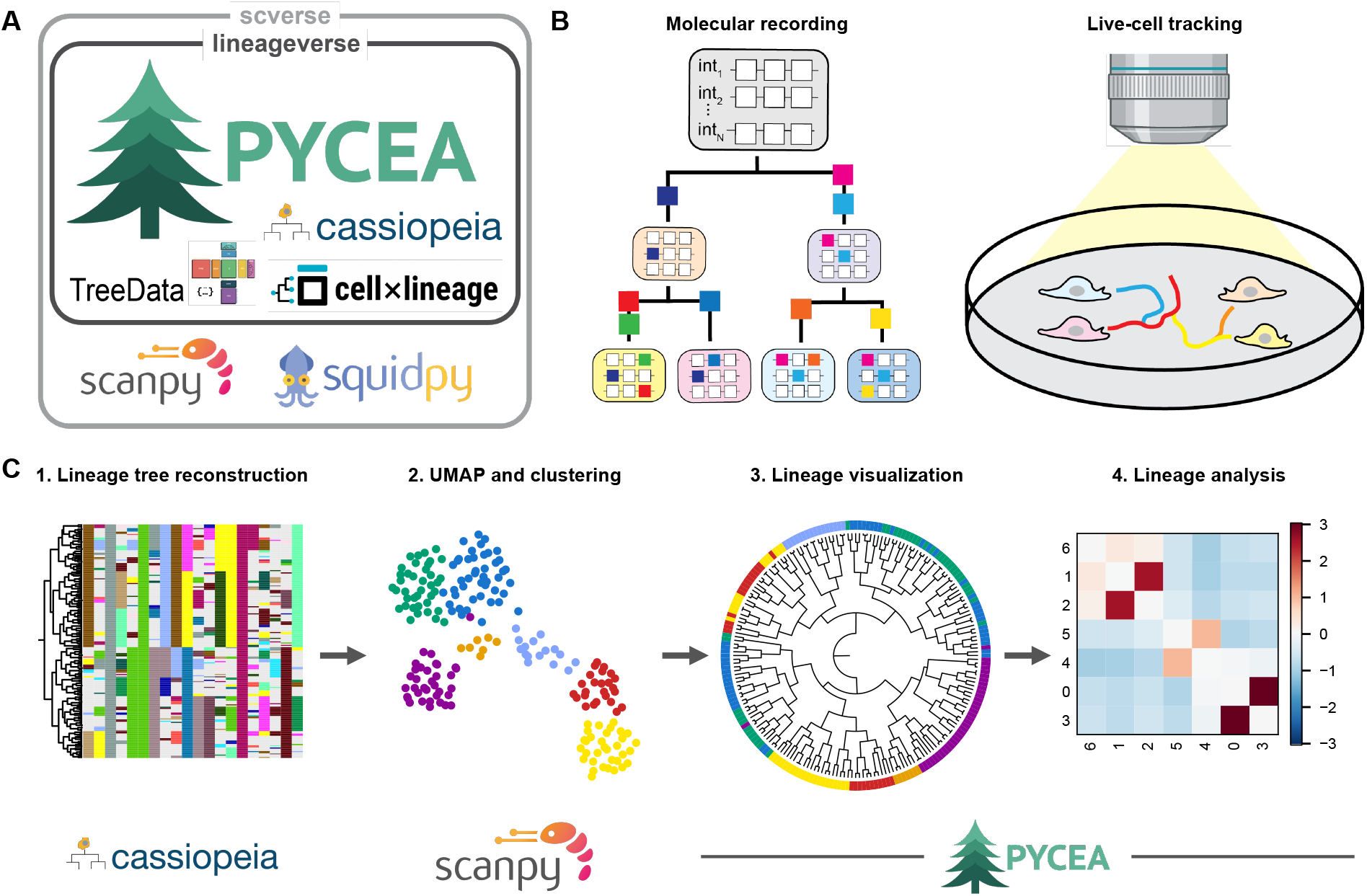
Pycea integrates lineage tracing analysis into the scverse ecosystem. **(A)** Pycea and TreeData within the scverse and lineageverse ecosystems. **(B)** Supported lineage tracing data: molecular recording (left) and live-cell tracking (right). **(C)** Example workflow on single-cell lineage tracing data: (1) lineage tree reconstruction with Cassiopeia, (2) UMAP and Leiden clustering with Scanpy, (3) polar tree with leaves annotated by cluster, and (4) ancestral linkage between clusters with Pycea.

## Results

### TreeData stores lineage trees alongside other single-cell measurements

To integrate lineage trees into standard single-cell workflows, we developed TreeData, an extension of AnnData that adds support for storing trees (Figure 2A). TreeData adds two fields, obst and vart, which store trees as `networkx` directed graphs [34] aligned with observations (cells) and variables (genes), respectively. Node and edge attributes, such as node time, are stored as key-value pairs on the tree. The AnnData interface (`X, obs, var, obsm, varm, layers, uns`, etc.) remains unchanged, and because TreeData is a subclass of AnnData, it can be passed directly to Scanpy, Squidpy, and other scverse tools. For molecular recording, the `obsm` field can store the character matrix used to reconstruct the tree, while for live-cell tracking, the obsm field can store cell coordinates and other measurements.

**Figure 2:**
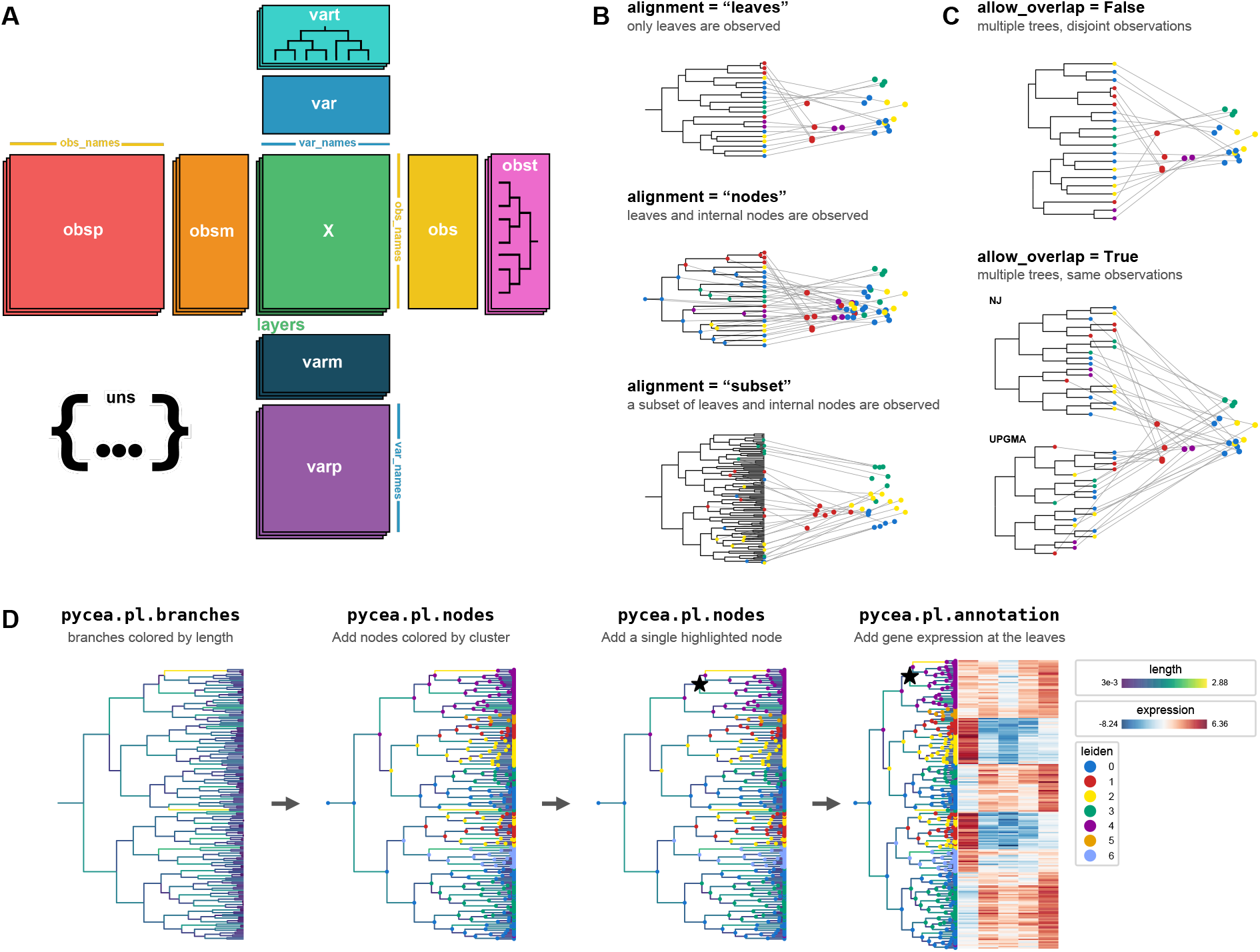
The TreeData object stores one or more trees aligned to observations. **(A)** TreeData extends AnnData with `obst` and `vart` fields, which store trees aligned to observations and variables. **(B)** Alignment modes. Gray lines connect tree nodes to their observations in the UMAP. With `“leaves”`, only leaves are observations; with `“nodes”`, leaves and internal nodes are; with “subset”, only some nodes are. **(C)** Multiple trees per object: trees with disjoint observations (`allow_overlap=False`, top) or multiple trees for the same observations (`allow_ overlap=True`, bottom; neighbor-joining (NJ) and UPGMA). **(D)** A tree plot built up with successive Pycea calls: pycea.pl.branches (branches colored by length), `pycea.pl.nodes` (leaves and internal nodes colored by cluster; then a single node highlighted), and `pycea.pl.annotation` (expression of five genes at the leaves).

TreeData objects can be created from scratch by specifying their constituent components or by converting an existing AnnData object. Trees stored in`obst` and `vart` can be assigned, modified, and deleted like any other field. When a TreeData object is sliced by observations or variables, the corresponding trees are automatically pruned to maintain consistency with the remaining data. TreeData objects can also be saved to disk as `.h5td` or `.zarr` files, with I/O times comparable to AnnData, particularly for scRNA-seq data where expression matrices dominate I/O (Figure S1A).

TreeData supports three `alignment` options (Figure 2B): (1) `“leaves”` indicates that observations correspond to the leaves of the tree, as is typically the case for molecular recording; (2) `“nodes”` indicates that both leaves and internal nodes correspond to observations, as is typically the case for live-cell tracking; and (3) `“subset”` indicates that only a subset of nodes correspond to observations (see `pycea.datasets.packer19` for an example in which a subset of nodes in the *C. elegans* lineage tree was measured [35]). Trees are validated against the observations upon assignment, and the selected `alignment` option controls how trees are pruned when the object is sliced. This pruning makes slicing noticeably slower than in AnnData (Figure S1A).

A TreeData object can hold multiple trees, each specified by a key (Figure 2C). For example, a TreeData object can store multiple trees from different tumor clones, mapped to a disjoint set of observations (`allow_overlap=False`, the default). Alternatively, multiple trees can describe the same observations (`allow_overlap=True`), as when comparing alternative reconstructions of the same lineage data (e.g. neighbor-joining and UPGMA trees). Pycea functions operate on all trees by default or on a selected subset. Thus, multi-clone experiments and comparisons of reconstruction methods do not require separate objects.

### Pycea provides composable tools for visualizing and analyzing lineage trees

Pycea is organized like Scanpy, with preprocessing (`pp`), tools (`tl`), plotting (`pl`) and data access (`get`) functions that operate on TreeData objects and write their results back to `obs, obsp, obst`, or `uns`. Pycea also provides a `datasets` module for loading example datasets, including the mouse embryo and tumor datasets analyzed below.

Complex tree plots can be constructed through successive `pl` calls (Figure 2D). For example, users can first draw tree branches with `pycea.pl.branches`, add nodes with `pycea.pl.nodes`, and finally add leaf annotations with `pycea.pl.annotation`. Each function can color the tree by attributes such as branch length, node depth, categorical or continuous annotations, and gene expression. Trees can be drawn in rectangular or polar layouts, and the plotting functions automatically add legends for categorical variables and colorbars for continuous variables. These functions scale to large trees, rendering trees with 100,000 cells in seconds (Figure S1B).

Pycea provides a suite of methods for quantifying lineage structure and relating it to other cellular measurements (Table S1). Distances and neighbor graphs derived from lineage trees, lineage barcode profiles, or other features are stored as sparse matrices in `obsp`, following Scanpy conventions, allowing them to be compared and used interchangeably in downstream analyses [36]. Tree topology and branch lengths can also be used to identify clades of closely related cells and estimate their relative fitness [37]. Other methods relate lineage structure to cellular phenotypes, including `ancestral_linkage`, which quantifies relatedness between cell populations, and `autocorr`, which measures the similarity of cellular attributes among relatives using Moran’s I [38]. Pycea also supports ancestral state reconstruction, inferring the states of internal nodes from observed cellular attributes using averaging or parsimony algorithms [39–41]. These methods scale to trees of 100,000 cells, with runtimes of 1–20 seconds for representative operations (Figure S1C), enabling integrated analysis of lineage, expression, and spatial data.

### Lineage relationships and gene heritability in a mouse embryo

To illustrate these tools, we analyzed mouse embryo lineage tracing data from Colgan et al. (`pycea.datasets.colgan26`) [2]. We focused on a single embryo collected at E9.5 (E9.5-R1), containing 262,534 lineage-traced cells. Each cell was assigned to a germ layer and a broad lineage category, such as neural crest, neural ectoderm, or surface ectoderm (Figure 3A–C). The lineage tree contains hundreds of distinct germ layer clades, reflecting substantial proliferation before germ layer specification.

**Figure 3:**
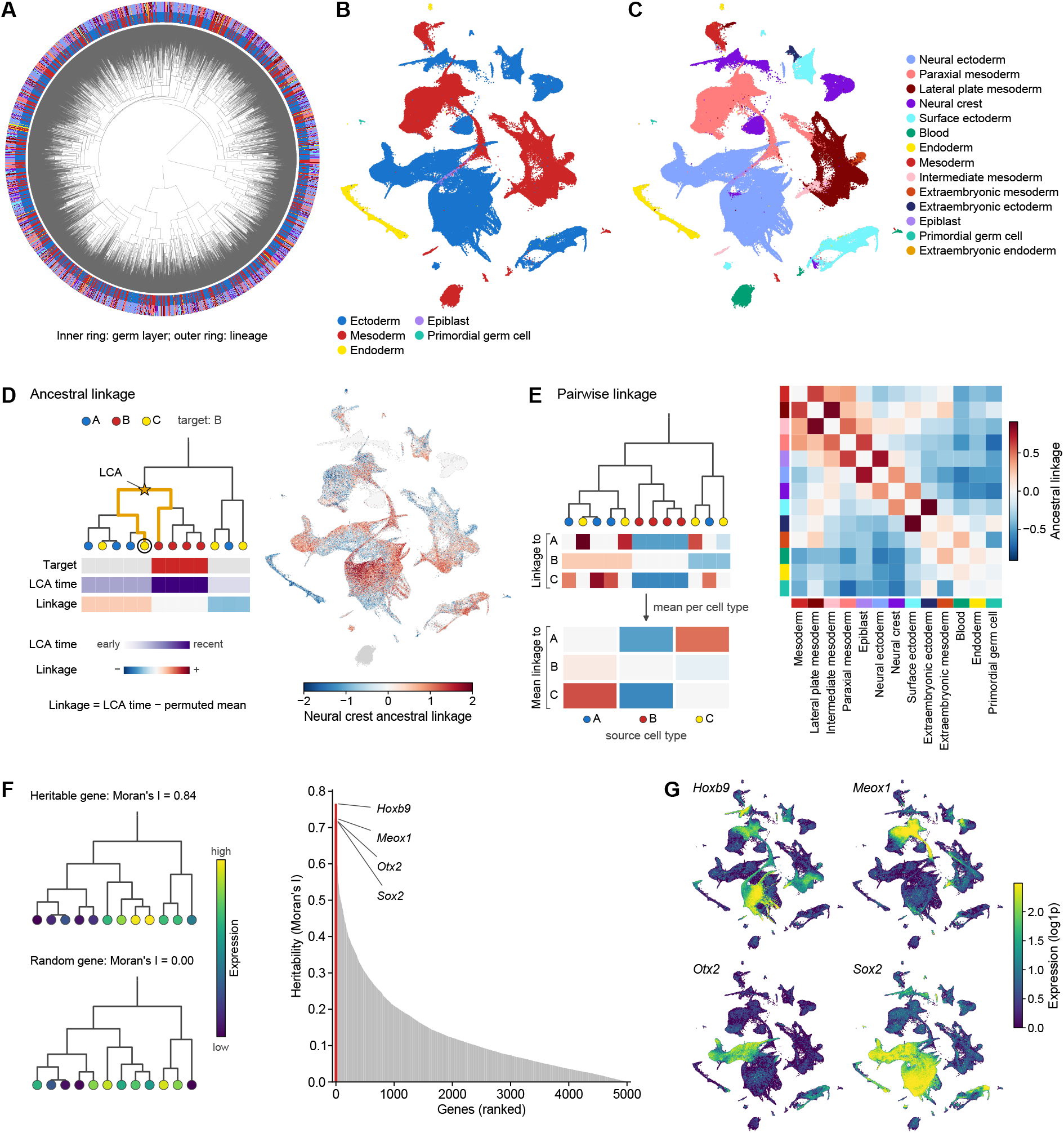
Lineage relationships and gene heritability in a mouse embryo. **(A)** Lineage trees of an E9.5 embryo (262,534 cells) with leaves annotated by germ layer (inner ring) and lineage (outer ring). **(B, C)** UMAP of cells colored by germ layer (B) and broad lineage annotation (C). **(D)** Ancestral linkage of a cell to a target set is the time of its most recent common ancestor (LCA) with the nearest target cell, normalized to chance (left). UMAP colored by neural crest ancestral linkage (right); gray cells are not in a lineage tree. **(E)** Pairwise linkage: the mean linkage of each cell type to a target cell type forms one row, repeated with every cell type as the target (left), giving the symmetrized linkage between all lineages (right; colors as in C). **(F)** Heritable genes have expression that is autocorrelated across the lineage tree, measured by Moran’s I (left), shown ranked for the 5,000 highly variable genes (right). **(G)** UMAPs colored by expression of the four most heritable genes.

We used `pycea.tl.ancestral_linkage` to examine the lineage relationships between different cell populations. For each cell, this function identifies the lowest common ancestor (LCA) shared with a target population (Figure 3D). Because this measure depends on tree topology and target population size, it is normalized by subtracting the mean under random permutation of the target cells (Methods) [10, 42]. For example, computing the ancestral linkage of every cell to the neural crest reveals that dorsal neural ectoderm and surface ectoderm cells share recent ancestry with this population (Figure 3D). We also computed mean pairwise linkage between lineages by averaging values for each source–target pair and symmetrizing the resulting matrix (Figure 3E). This analysis groups lineages by germ layer and reveals a notable association between the neural crest and intermediate mesoderm, potentially reflecting their similar mediolateral positions in the embryo. Pycea also enables gene-level analysis on trees. For example, we quantified the heritability of gene expression across the lineage tree using Moran’s I autocorrelation statistic (`pycea.tl. autocorr`) [38]. Genes whose expression is unrelated to lineage have values near 0, whereas genes with correlated expression among closely related cells have higher values (Figure 3F). In the mouse embryo, the four most heritable genes were *Hoxb9, Meox1, Otx2* and *Sox2*, all of which are lineage-defining transcription factors or patterning genes whose expression is restricted to specific regions of the UMAP (Figure 3G).

### Cell proliferation dynamics in a tumor metastasis model

To demonstrate how Pycea can be used to analyze spatially resolved lineage tracing data, we analyzed a 4T1 tumor dataset from Koblan et al. (`pycea.datasets.koblan25`) [15]. We focused on three sections from a single tumor (M3-T1), containing 145,954 cells with gene expression and lineage information measured *in situ* using imaging. We used `pycea.tl.clades` to group cells into clades based on shared ancestry at day 5, inferred from estimated branch lengths, and plotted these clades alongside the lineage marks used to reconstruct the tree (Figure 4A). Because TreeData is compatible with Squidpy, we could readily visualize the spatial distribution of these clades, revealing that spatially proximal cells are more likely to share recent ancestry (Figure 4B).

**Figure 4:**
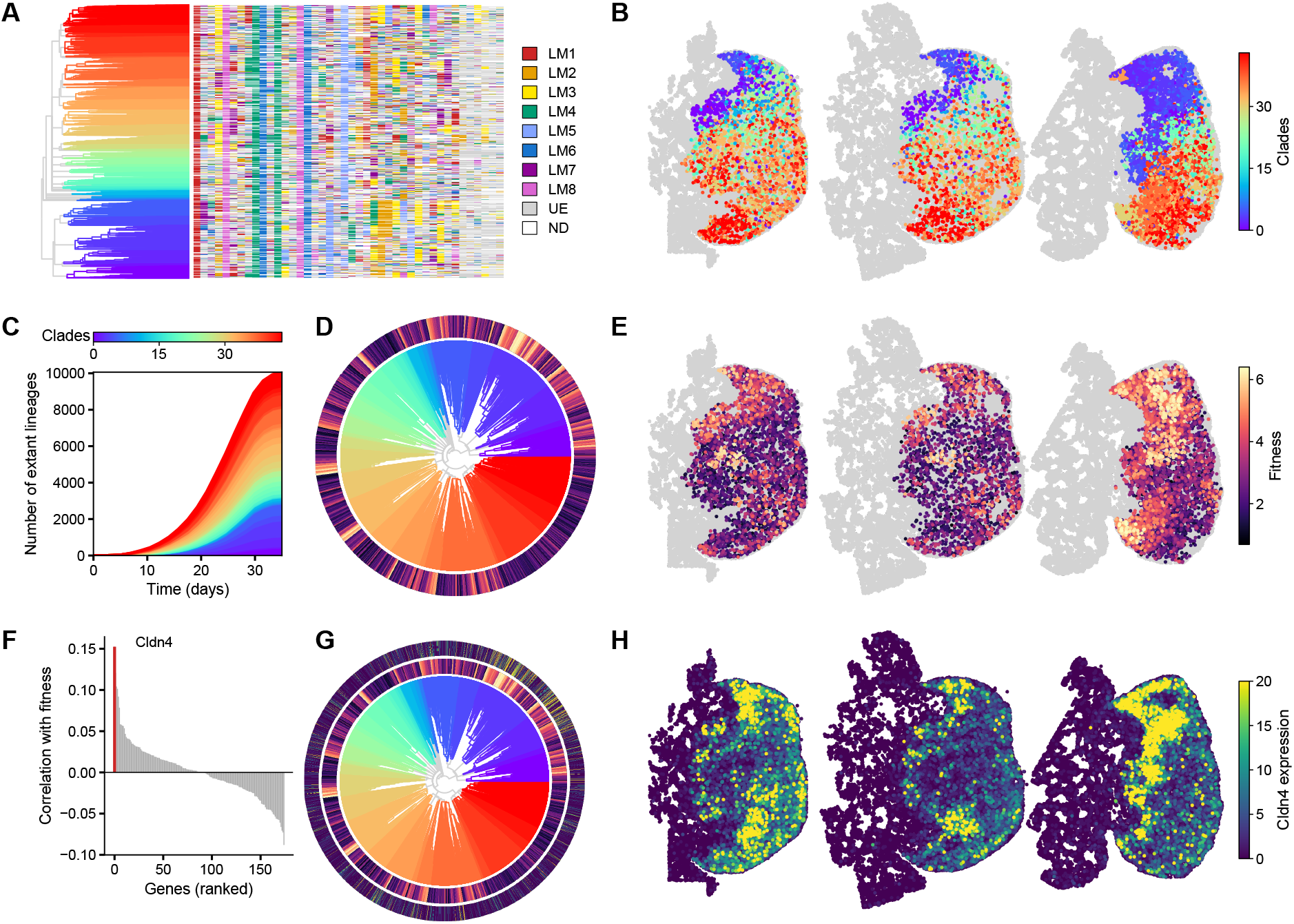
Growth dynamics of a 4T1 tumor. **(A)** Lineage tree and character matrix, with branches colored by clade (cells sharing an ancestor at day 5); colors indicate lineage marks (LMs), gray unedited (UE), and white no detection (ND). **(B)** Spatial distribution of clades across three tumor sections; gray cells are not in the lineage tree. **(C)** Number of extant lineages in each clade over time (`pycea.pl.n_extant`). **(D)** Polar tree with the estimated fitness (`pycea.tl.fitness`) of each cell in the outer ring. **(E)** Spatial distribution of fitness. **(F)** Correlation of gene expression with fitness; *Cldn4* is highlighted. **(G)** Polar tree with rings for fitness and *Cldn4* expression. **(H)** Spatial distribution of *Cldn4* expression.

We then used the tree to quantify proliferation dynamics. `pycea.tl.n_extant` counts the lineages alive at each time point, revealing how the relative abundance of these clades changes over time (Figure 4C). `pycea.tl.fitness` estimates the relative fitness of each cell from tree topology and branch lengths using selection-biased diffusion [37] (Figure 4D,E; Methods). Notably, the blue and purple clades tended to have higher fitness, with the highest-fitness cells occupying the tumor periphery near normal lung tissue. We also investigated whether gene expression was associated with fitness by computing the Pearson correlation between each gene’s expression and fitness across cells (Figure 4F). The strongest correlation was observed for *Cldn4*, which is expressed in the lung-adjacent tumor and associated with an epithelial-like state (Figure 4G,H) [15].

## Discussion

We introduce TreeData as a standardized data structure for lineage trees within the scverse ecosystem and Pycea as a toolkit for lineage analysis and visualization. Together, they enable lineage, gene expression, and spatial data to be analyzed within a unified framework, simplifying workflows that previously required multiple data structures and tools. Pycea and [TreeData](https://github.com/YosefLab/treedata) are open source, with documentation and tutorials reproducing the analyses presented here [available online](https://pycea.readthedocs.io/).

We envision TreeData serving as the foundation for a growing ecosystem of interoperable lineage analysis tools, which we refer to as the ‘lineageverse’. As a step toward this goal, the upcoming Cassiopeia v3.0.0 release will replace the existing CassiopeiaTree object with TreeData. This will allow users to seamlessly combine Cassiopeia for preprocessing lineage data and reconstructing trees, Scanpy for transcriptomic analysis, and Pycea for lineage analysis and visualization. We will continue to expand this ecosystem with additional TreeData-compatible tools, such as the [CELLxLINEAGE viewer](https://github.com/colganwi/cellxlineage), and encourage the community to adopt and extend this framework.

## Methods

### Software

TreeData (v0.3.1) extends the AnnData object with `obst` and `vart` fields, which store lineage trees as `networkx` directed graphs [34] aligned to the observations and variables. Pycea (v0.4.0) provides tools (`pycea.tl`), plotting (`pycea.pl`), preprocessing (`pycea.pp`), and data retrieval (`pycea.get`) modules that operate on TreeData objects. Analyses used Python 3.12, Scanpy [24] 1.12 and AnnData 0.13. Trees are plotted with `pycea.pl.branches, pycea.pl.nodes` and `pycea.pl. annotation`, which draw the branches, nodes and leaf annotations of a tree in rectangular or polar coordinates.

### Datasets

#### Simulated data

The lineage trees, characters and gene expression in Figures 1 and 2 were simulated with Cassiopeia [32] (v3.0.0). Neighbor-joining and UPGMA trees were reconstructed from the simulated characters with `cassiopeia.solver.nj` and `cassiopeia.solver.upgma.`

#### Mouse embryo

We analyzed embryo E9.5-R1 from the PEtracer mouse embryo atlas [2]. Annotations, UMAP coordinates and trees were loaded with `pycea.datasets.colgan26`. Gene expression counts were normalized to 10,000 counts per cell and log1p-transformed, and the 5,000 highly variable genes defined in the atlas were analyzed.

#### Tumor

We analyzed spatially resolved lineage tracing of 4T1 tumors [15] (M3-T1), loaded with `pycea.datasets.koblan25`. Analyses were restricted to the three tumor sections that contain lineage-traced cells.

### Ancestral linkage

`pycea.tl.ancestral_linkage` quantifies how closely related cells are to a target population. It supports two types of analysis: (1) measuring the ancestral linkage of individual cells to a target population, and (2) comparing the average ancestral linkage between pairs of populations. For example, we can measure how closely each cell in an embryo is related to the neural crest (Figure 3D), or compare the ancestral relationships between all major cell lineages (Figure 3E).

For a cell *i* and target category *T*, linkage is defined as the depth of its most recent common ancestor with any cell in *T*,

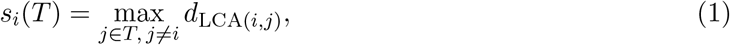

where *d*_LCA(*i,j*)_ is the depth (time) of the lowest common ancestor of cells *i* and *j*, with larger values indicating more recent common ancestry. Because tree topology and category size affect this value, scores are normalized by subtracting their mean under random permutation of the category labels, 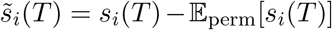. Thus, 0 represents the chance expectation, and positive values indicate greater relatedness to *T* than expected. Per-cell linkage to the neural crest (Figure 3D) was computed with `groupby=“lineage” and target=“Neural crest”.`

To compare pairs of populations, the normalized linkage scores are averaged over all cells in each source category *a* relative to a target category *b*,

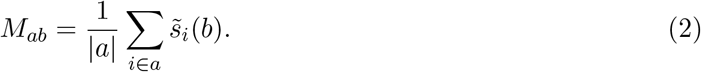

The resulting directional matrix is symmetrized by averaging (*M*_*ab*_ + *M*_*ba*_)*/*2. Pairwise linkage between lineages was computed with `target=None` and visualized using `pycea.pl.ancestral_ linkage` (Figure 3E).

### Gene heritability

Heritability of gene expression was estimated as its autocorrelation over the lineage tree. `pycea. tl.tree_neighbors` (`n_neighbors=5, depth_key=“time”`) identified the five nearest relatives of each cell by tree path distance, *d*_*i*_ + *d*_*j*_ − 2 *d*_LCA(*i,j*)_, and stored them as a sparse connectivity matrix with weights *w*_*ij*_. `pycea.tl.autocorr` (`method=“moran”`) then computed Moran’s I for each gene,

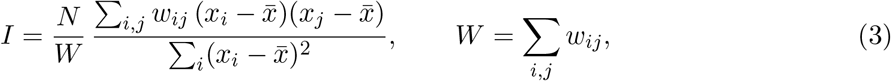

where *x*_*i*_ is the expression (log1p) of the gene in cell *i*, 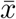its mean and *N* the number of cells. Values near 1 indicate that closely related cells have similar expression (heritable), and values near 0 indicate no relationship to the lineage. Genes were ranked by *I* (Figure 3F) and the four highest are shown (Figure 3G).

### Clades and fitness

Clades (Figure 4A, B) were defined with `pycea.tl.clades` (`depth=5, depth_key=“time”`): each branch alive at day 5 defines a clade, and all of its descendants inherit its label. The number of extant lineages in each clade over time (Figure 4C) was computed with `pycea.tl.n_extant` as the number of branches spanning each of 20 time points, grouped by clade, and plotted with `pycea.pl. n_extant.` Relative fitness of each cell (Figure 4D, E) was estimated from the tree topology and branch lengths with `pycea.tl.fitness` (`method=“sbd”`), which implements the selection-biased diffusion algorithm of Neher et al. [37]: messages are passed up and down the tree to infer the posterior distribution of fitness at each node, and the posterior mean is reported (diffusion constant *γ* = 0.2). Genes associated with fitness (Figure 4F) were identified by the Pearson correlation between the fitness and expression of each gene across cells with both measurements; *Cldn4*, the most positively correlated gene, is shown alongside fitness on the tree and in space (Figure 4G, H). Spatial panels (Figure 4B, E, H) plot the measured cell coordinates with Scanpy.

### Benchmarks

Lineage trees of 10^2^ to 10^5^ cells were simulated with Cassiopeia (v3.0.0). TreeData and AnnData objects with simulated sparse expression of 20,000 genes (5% nonzero entries) were written to and read from disk (`.h5td` and `.h5ad`), and sliced to a random 10% of cells, which includes pruning the tree and materializing a copy. Tree plotting time (`pycea.pl.tree`, rectangular layout) was compared with Cassiopeia 2.1.0 (`cassiopeia.pl.plot_matplotlib`) and includes rendering the figure canvas. The run time of `pycea.pp.add_depth, pycea.tl.clades, pycea.tl.ancestral_ states, pycea.tl.tree_neighbors` (10 neighbors), `pycea.tl.ancestral_linkage`, `pycea.tl. fitness` (`method=“lbi”`) and `pycea.tl.expansion_test` was measured on trees with five random categories. Each measurement is the median of three runs (a single run if the first took longer than 30 s), on a single Slurm node with two CPU cores.

### Use of large language models

Claude and ChatGPT were used to assist with writing, analysis, and code generation. The authors carefully review all model outputs and take full responsibility for the content of the manuscript.

## Declarations

### Ethics approval and consent to participate

Not applicable.

### Consent for publication

Not applicable.

### Availability of data and materials

Pycea and [TreeData](https://github.com/YosefLab/treedata) are open source under the BSD 3-Clause license and are available on GitHub and PyPI (`pycea-lineage and treedata`), with documentation and tutorials. The versions described here are Pycea v0.4.0 and TreeData v0.3.1. Code to reproduce all figures is available at https://github.com/colganwi/pycea-paper. The data for Figures 1, 2 and S1 are simulated by scripts in that repository.

The mouse embryo data (E9.5-R1) are from Colgan et al. [2] and can be downloaded from Zenodo. The 4T1 tumor data are from Koblan et al. [15] and can be downloaded from Figshare. Processed versions of both datasets, which are loaded by `pycea.datasets.colgan26` and `pycea. datasets.koblan25`, are available on Zenodo.

### Authors’ contributions

Conceptualization: WNC, LWK, MGJ, JSW, NY

Methodology: WNC, MGJ

Software: WNC, EW, MGJ

Formal analysis: WNC

Resources: JSW, NY

Data curation: WNC

Writing – original draft: WNC

Writing – review and editing: WNC, EW, LWK, MGJ, JSW, NY

Visualization: WNC, LWK

Supervision: JSW, NY

Funding acquisition: JSW, NY

All authors read and approved the final manuscript.

## Acknowledgements

We thank Sebastian Prillo, Pedro Olivares, Yaara Neumeier, and Dana Rimini for helpful discussions.

---

**Figure S1:**
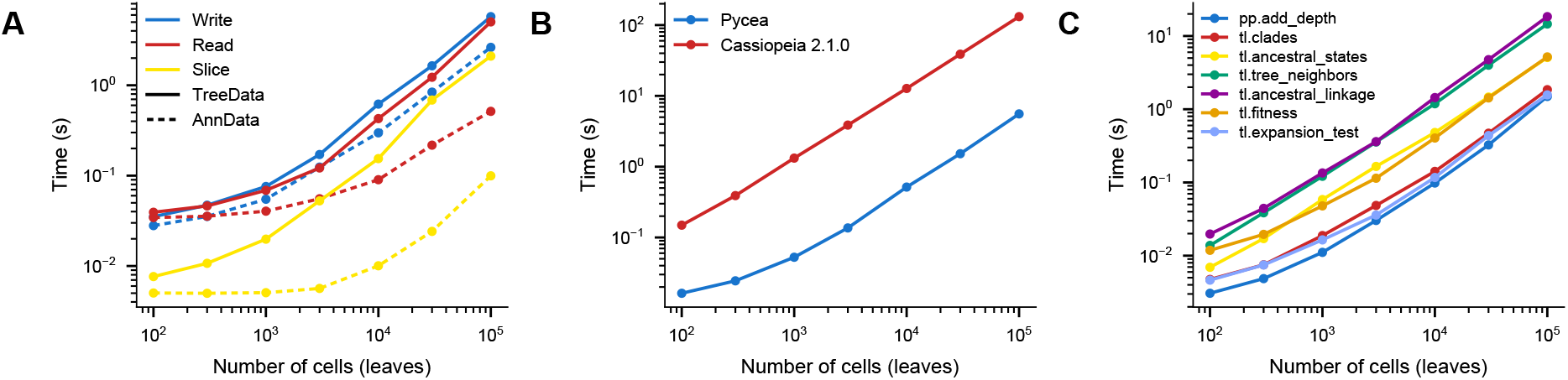
Benchmarks. **(A)** Time to write, read and slice (a random 10% of cells) TreeData and AnnData objects with a simulated lineage tree and simulated expression of 20,000 genes (sparse counts), by number of cells. **(B)** Time to plot a tree with Pycea and with Cassiopeia 2.1.0 (plotting deprecated in version 3.0.0), by number of cells. **(C)** Run time of key Pycea functions, by number of cells. Median of three runs, on a single node.

**Table S1:** Public functions of Pycea v0.4.0. The 33 functions in the Pycea modules. See the Pycea API documentation for details.

| Function | Description |
|---|---|
| **pycea.pp**: Preprocessing |  |
| **add_depth** | Add a depth attribute (e.g., time) to each node of the tree |
| **pycea.tl**: Tools |  |
| **ancestral_linkage** | Measure relatedness between categories of cells |
| **ancestral_states** | Reconstruct internal node states (e.g., parsimony [39–41]) |
| **autocorr** | Calculate Moran's I or Geary's C over a neighbor graph [38] |
| **clades** | Assign cells and nodes to clades |
| **compare_distance** | Compare distances between the same pairs of cells |
| **distance** | Compute distances between cells in any embedding |
| **expansion_test** | Test subclades for expansion under neutral growth [43] |
| **fitch_count** | Count transitions between states [44] |
| **fitness** | Estimate the relative fitness of each node [37] |
| **n_extant** | Count extant lineages over time |
| **neighbor_distance** | Aggregate distances from each cell to its neighbors |
| **parsimony** | Compute the small-parsimony score of a tree |
| **partition_test** | Compare the leaves below a node with its siblings or the rest |
| **sort** | Reorder branches by a node attribute |
| **tree_distance** | Compute distances between cells in the tree |
| **tree_neighbors** | Identify the closest relatives of each cell in the tree |
| **pycea.pl**: Plotting |  |
| **ancestral_linkage** | Plot ancestral linkage as a clustered heatmap |
| **annotation** | Plot leaf annotations (e.g., metadata, expression) beside a tree |
| **branches** | Plot the branches of a tree |
| **n_extant** | Plot the number of extant lineages over time |
| **nodes** | Plot the nodes of a tree |
| **tree** | Plot a tree with branches, nodes, and annotations |
| **pycea.get**: Data access |  |
| **edge_df** | Get tree edge attributes as a data frame |
| **leaves** | Get the leaves of a tree |
| **node_df** | Get tree node attributes as a data frame |
| **palette** | Get the color palette for a categorical variable |
| **root** | Get the root of a tree |
| **pycea.datasets**: Example datasets |  |
| **colgan26** | Mouse embryo lineage tracing, E7.5–E10.0 [2] |
| **koblan25** | Spatial lineage tracing of 4T1 tumors [15] |
| **packer19** | *C. elegans* lineage tree and cell states [35] |
| **yang22** | KP mouse model lineage tracing [43] |
| **yu26** | DNA Typewriter mouse lineage tracing [1] |

## References

[1] Yu Q, Kim H, Seidel S, Acosta-Clark JF, Martin BK, O’Connor K, et al. A DNA Type-writer records the cell lineage history of a mouse, from zygote to late organogenesis. Science. 2026:eael0508. doi:10.1126/science.ael0508.

[2] Colgan WN, Koblan LW, Villagrana J, Hou TCJ, Wang M, Gowri G, et al. Comprehen-sive lineage tracing maps the landscape of cell fate decisions in mouse embryogenesis. Cell. 2026;189:1–23. doi:10.1016/j.cell.2026.09.050.

[3] Sulston JE, Schierenberg E, White JG, Thomson JN. The embryonic cell lineage of the nematode Caenorhabditis elegans. Dev Biol. 1983;100(1):64–119. doi:10.1016/0012-1606(83)90201-4.

[4] Keller PJ, Schmidt AD, Wittbrodt J, Stelzer EHK. Reconstruction of Zebrafish Early Embryonic Development by Scanned Light Sheet Microscopy. Science. 2008;322(5904):1065–9. doi:10.1126/science.1162493.

[5] McDole K, Guignard L, Amat F, Berger A, Malandain G, Royer LA, et al. In Toto Imaging and Reconstruction of Post-Implantation Mouse Development at the Single-Cell Level. Cell. 2018;175(3):859-76.e33. doi:10.1016/j.cell.2018.09.031.

[6] Ulman V, Maška M, Magnusson KEG, Ronneberger O, Haubold C, Harder N, et al. An objective comparison of cell-tracking algorithms. Nat Methods. 2017;14(12):1141–52. doi:10.1038/nmeth.4473.

[7] Ershov D, Phan MS, Pylvänäinen JW, Rigaud SU, Le Blanc L, Charles-Orszag A, et al. TrackMate 7: integrating state-of-the-art segmentation algorithms into tracking pipelines. Nat Methods. 2022;19(7):829–32. doi:10.1038/s41592-022-01507-1.

[8] McKenna A, Findlay GM, Gagnon JA, Horwitz MS, Schier AF, Shendure J. Wholeorganism lineage tracing by combinatorial and cumulative genome editing. Science. 2016;353(6298):aaf7907. doi:10.1126/science.aaf7907.

[9] Kalhor R, Kalhor K, Mejia L, Leeper K, Graveline A, Mali P, et al. Developmental barcoding of whole mouse via homing CRISPR. Science. 2018;361(6405):eaat9804. doi:10.1126/science.aat9804.

[10] Chan MM, Smith ZD, Grosswendt S, Kretzmer H, Norman TM, Adamson B, et al. Molecular recording of mammalian embryogenesis. Nature. 2019;570(7759):77–82. doi:10.1038/s41586-019-1184-5.

[11] Bowling S, Sritharan D, Osorio FG, Nguyen M, Cheung P, Rodriguez-Fraticelli A, et al. An Engineered CRISPR-Cas9 Mouse Line for Simultaneous Readout of Lineage Histories and Gene Expression Profiles in Single Cells. Cell. 2020;181(6):1410-22.e27. doi:10.1016/j.cell.2020.04.048.

[12] Frieda KL, Linton JM, Hormoz S, Choi J, Chow KHK, Singer ZS, et al. Synthetic recording and in situ readout of lineage information in single cells. Nature. 2017;541(7635):107–11. doi:10.1038/nature20777.

[13] Chow KHK, Budde MW, Granados AA, Cabrera M, Yoon S, Cho S, et al. Imaging cell lineage with a synthetic digital recording system. Science. 2021;372(6538):eabb3099. doi:10.1126/science.abb3099.

[14] Choi J, Chen W, Minkina A, Chardon FM, Suiter CC, Regalado SG, et al. A time-resolved, multi-symbol molecular recorder via sequential genome editing. Nature. 2022;608(7921):98–107. doi:10.1038/s41586-022-04922-8.

[15] Koblan LW, Yost KE, Zheng P, Colgan WN, Jones MG, Yang D, et al. High-resolution spatial mapping of cell state and lineage dynamics in vivo with PEtracer. Science. 2025;390(6770):eadx3800. doi:10.1126/science.adx3800.

[16] Hwang B, Lee W, Yum SY, Jeon Y, Cho N, Jang G, et al. Lineage tracing using a Cas9-deaminase barcoding system targeting endogenous L1 elements. Nat Commun. 2019;10(1):1234. doi:10.1038/s41467-019-09203-z.

[17] Winter E, Emiliani F, Cook A, Abderrahim A, McKenna A. BASELINE: a CRISPR base editing platform for mammalian-scale single-cell lineage tracing. Nucleic Acids Res. 2026;54(14):gkag769. doi:10.1093/nar/gkag769.

[18] Raj B, Wagner DE, McKenna A, Pandey S, Klein AM, Shendure J, et al. Simultaneous single-cell profiling of lineages and cell types in the vertebrate brain. Nat Biotechnol. 2018;36(5):442–50. doi:10.1038/nbt.4103.

[19] Spanjaard B, Hu B, Mitic N, Olivares-Chauvet P, Janjuha S, Ninov N, et al. Simultaneous lineage tracing and cell-type identification using CRISPR–Cas9-induced genetic scars. Nat Biotechnol. 2018;36(5):469–73. doi:10.1038/nbt.4124.

[20] Virshup I, Bredikhin D, Heumos L, Palla G, Sturm G, Gayoso A, et al. The scverse project provides a computational ecosystem for single-cell omics data analysis. Nat Biotechnol. 2023;41(5):604–6. doi:10.1038/s41587-023-01733-8.

[21] Virshup I, Rybakov S, Theis FJ, Angerer P, Wolf FA. anndata: Access and store annotated data matrices. J Open Source Softw. 2024;9(101):4371. doi:10.21105/joss.04371.

[22] Bredikhin D, Kats I, Stegle O. MUON: multimodal omics analysis framework. Genome Biol. 2022;23(1):42. doi:10.1186/s13059-021-02577-8.

[23] Marconato L, Palla G, Yamauchi KA, Virshup I, Heidari E, Treis T, et al. SpatialData: an open and universal data framework for spatial omics. Nat Methods. 2025;22(1):58–62. doi:10.1038/s41592-024-02212-x.

[24] Wolf FA, Angerer P, Theis FJ. SCANPY: large-scale single-cell gene expression data analysis. Genome Biol. 2018;19(1):15. doi:10.1186/s13059-017-1382-0.

[25] Palla G, Spitzer H, Klein M, Fischer D, Schaar AC, Kuemmerle LB, et al. Squidpy: a scalable framework for spatial omics analysis. Nat Methods. 2022;19(2):171–8. doi:10.1038/s41592-021-01358-2.

[26] Zafar H, Lin C, Bar-Joseph Z. Single-cell lineage tracing by integrating CRISPR-Cas9 muttions with transcriptomic data. Nat Commun. 2020;11(1):3055. doi:10.1038/s41467-020-16821-5.

[27] Gong W, Kim HJ, Garry DJ, Kwak IY. Single cell lineage reconstruction using distance-based algorithms and the R package, DCLEAR. BMC Bioinformatics. 2022;23(1):103. doi:10.1186/s12859-022-04633-x.

[28] Sashittal P, Schmidt H, Chan M, Raphael BJ. Startle: A star homoplasy approach for CRISPR-Cas9 lineage tracing. Cell Syst. 2023;14(12):1113-21.e9. doi:10.1016/j.cels.2023.11.005.

[29] Seidel S, Stadler T. TiDeTree: a Bayesian phylogenetic framework to estimate single-cell trees and population dynamic parameters from genetic lineage tracing data. Proc Biol Sci. 2022;289(1986):20221844. doi:10.1098/rspb.2022.1844.

[30] Chu G, Mai U, Schmidt H, Raphael BJ. Maximum likelihood inference of time-scaled cell lineage trees with mixed-type missing data using LAML. Genome Biol. 2025;26(1):189. doi:10.1186/s13059-025-03649-9.

[31] Chu G, Schmidt H, Raphael BJ. LAML-Pro: joint maximum likelihood inference of cell genotypes and cell lineage trees. Bioinformatics. 2026;42:btag219. doi:10.1093/bioinformatics/btag219.

[32] Jones MG, Khodaverdian A, Quinn JJ, Chan MM, Hussmann JA, Wang R, et al. Inference of single-cell phylogenies from lineage tracing data using Cassiopeia. Genome Biol. 2020;21(1):92. doi:10.1186/s13059-020-02000-8.

[33] Huerta-Cepas J, Serra F, Bork P. ETE 3: Reconstruction, Analysis, and Visualization of Phylogenomic Data. Mol Biol Evol. 2016;33(6):1635–8. doi:10.1093/molbev/msw046.

[34] Hagberg AA, Schult DA, Swart PJ. Exploring network structure, dynamics, and function using NetworkX. In: Varoquaux G, Vaught T, Millman J, editors. Proceedings of the 7th Python in Science Conference (SciPy2008). Pasadena, CA, USA; 2008. p. 11–5. doi:10.25080/tcwv9851.

[35] Packer JS, Zhu Q, Huynh C, Sivaramakrishnan P, Preston E, Dueck H, et al. A lineage-resolved molecular atlas of C. elegans embryogenesis at single-cell resolution. Science. 2019;365(6459):eaax1971. doi:10.1126/science.aax1971.

[36] DeTomaso D, Yosef N. Hotspot identifies informative gene modules across modalities of single-cell genomics. Cell Syst. 2021;12(5):446-56.e9. doi:10.1016/j.cels.2021.04.005.

[37] Neher RA, Russell CA, Shraiman BI. Predicting evolution from the shape of genealogical trees. eLife. 2014;3:e03568. doi:10.7554/eLife.03568.

[38] Moran PAP. Notes on continuous stochastic phenomena.

[39] Fitch WM. Toward Defining the Course of Evolution: Minimum Change for a Specific Tree Topology. Syst Zool. 1971;20(4):406–16. doi:10.2307/2412116.

[40] Hartigan JA. Minimum Mutation Fits to a Given Tree. Biometrics. 1973;29(1):53–65. doi:10.2307/2529676.

[41] Sankoff D. Minimal Mutation Trees of Sequences. SIAM J Appl Math. 1975;28(1):35–42. doi:10.1137/0128004.

[42] Fang W, Bell CM, Sapirstein A, Asami S, Leeper K, Zack DJ, et al. Quantitative fate mapping: A general framework for analyzing progenitor state dynamics via retrospective lineage barcoding. Cell. 2022;185(24):4604-20.e32. doi:10.1016/j.cell.2022.10.028.

[43] Yang D, Jones MG, Naranjo S, Rideout WM, Min KHJ, Ho R, et al. Lineage tracing reveals the phylodynamics, plasticity, and paths of tumor evolution. Cell. 2022;185(11):1905-23.e25. doi:10.1016/j.cell.2022.04.015.

[44] Quinn JJ, Jones MG, Okimoto RA, Nanjo S, Chan MM, Yosef N, et al. Single-cell lineages reveal the rates, routes, and drivers of metastasis in cancer xenografts. Science. 2021;371(6532):eabc1944. doi:10.1126/science.abc1944.

